# Complex spatio-temporal distribution and genogeographic affinity of mitochondrial DNA haplogroups in 24,216 Danes

**DOI:** 10.1101/148494

**Authors:** Jonas Bybjerg-Grauholm, Christian M Hagen, Vanessa F Gonçalves, Marie Bækvad-Hansen, Christine S Hansen, Paula L Hedley, Jørgen K Kanters, Jimmi Nielsen, Michael Theisen, Ole Mors, James Kennedy, Thomas D Als, Alfonso B Demur, Thomas M Werge, Merete Nordentoft, Anders Børglum, Preben Bo Mortensen, David M Hougaard, Michael Christiansen

**Affiliations:** Department for Congenital Disorders, Statens Serum Institut, Copenhagen, Denmark; Centre for Addiction and Mental Health, University of Toronto, Toronto, Canada; Department of Biomedical Sciences, University of Copenhagen, Copenhagen, Denmark; Aalborg Psychiatric Hospital. Aalborg University Hospital, Aalborg, Denmark; Department of Clinical Medicine, Aarhus University, Århus, Denmark; Institute of Medical Genetics, Aarhus University, Århus, Denmark; Mental Health Centre, Sct Hans, Capital Region of Denmark, Denmark; Mental Health Centre, Capital Region of Denmark, Denmark; Center for Register Research, Institute of Economics, Aarhus University, Århus, Denmark

**Author notes:** The study was conducted under the auspices of the iPSYCH study (www.iPSYCH.au.dk). JG and CMH contributed equally to the study. **Correspondence:** Professor, chief physician, Michael Christiansen, FRCPath, MD, Department for Congenital Disorders, Statens Serum Institut And Department of Biomedical Sciences, University of Copenhagen.

**Keywords:** mitochondrial DNA, haplogroup, population genetics, energy metabolism

## Abstract

Mitochondrial DNA (mtDNA) haplogroups (hgs) are evolutionarily conserved sets of mtDNA SNP-haplotypes with characteristic geographical distribution. Associations of hgs with disease and physiological characteristics have been reported, but have frequently not been reproducible. Using 418 mtDNA SNPs on the PsychChip (Illumina), we assessed the spatio-temporal distribution of mtDNA hgs in Denmark in DNA isolated from 24,642 geographically un-biased dried blood spots (DBS), collected from 1981 to 2005 through the Danish National Neonatal Screening program. Geno-geographic affinity (ancestry background) was established with ADMIXTURE using a reference of 100K+ autosomal SNPs in 2,248 individuals from nine populations. The hg distribution was typically Northern European, and hgs were highly variable based on median-joining analysis, suggesting multiple founder events. Considerable heterogeneity and variation in autosomal geno-geographic affinity was observed. Thus, individuals with hg H exhibited 95 %, and U hgs 38.2 % - 92.5 %, Danish ancestry. Significant clines between geographical regions and rural and metropolitan populations were found. Over 25 years, macro-hg L increased from 0.2 % to 1.2 % (p = 1.1*E-10), and M from 1 % to 2.4 % (p = 3.7*E-8). Hg U increased among the R macro-hg from 14.1 % to 16.5 % (p = 1.9*E-3). Geno-geographic affinity, geographical skewedness, and sub-hg distribution suggested that the L, M and U increases are due to immigration. The complex spatio-temporal dynamics and geno-geographic heterogeneity of mtDNA in the Danish population reflect repeated migratory events and, in later years, net immigration. Such complexity may explain the often contradictory and population-specific reports of mito-genomic association with disease.

## Introduction

Mitochondria are subcellular organelles responsible for oxidative phosphorylation (OXPHOS), producing ~ 80% of the ATP in eukaryotic cells^1^, apoptosis and cell-cycle regulation^2^, redox- and calcium homeostasis^3^ as well as intracellular signaling^4^. Each mitochondrion contains 2 - 10 copies of a 16.6 kb double-stranded mtDNA containing 37 genes^5^. Thirteen genes code for proteins in the five enzyme complexes conducting OXPHOS, whereas twenty-two genes code for tRNAs and two for rRNAs, all involved in intra-mitochondrial translation^6^. The mitochondrial proteome comprises approximately 1200 proteins^7^; ^8^, of which mtDNA genes encode ~ 1%. The mtDNA is maternally inherited^9^, exhibits a high mutation rate^10^, and does not undergo recombination^5^. Genetic variants in mtDNA – as well as variants in the nuclear genes encoding the mitochondrial proteome - have been associated with disease^11^; ^12^ ^13^. More than 150 mitochondrial syndromes^14^ have been associated with more than 300 variants^7^; ^15^.

Geographically and population specific lineages of mtDNA, haplogroups (hgs), have become fixed^17^, through the processes of random genetic drift and selection as the human populations dispersed throughout the world^16^. The advent of high throughput DNA sequencing technology, as well as implementation of biobanking technologies^18^, has enabled the construction of a high-resolution phylogenetic matrilineal mtDNA tree, Figure 1^19^.

**Figure 1.**
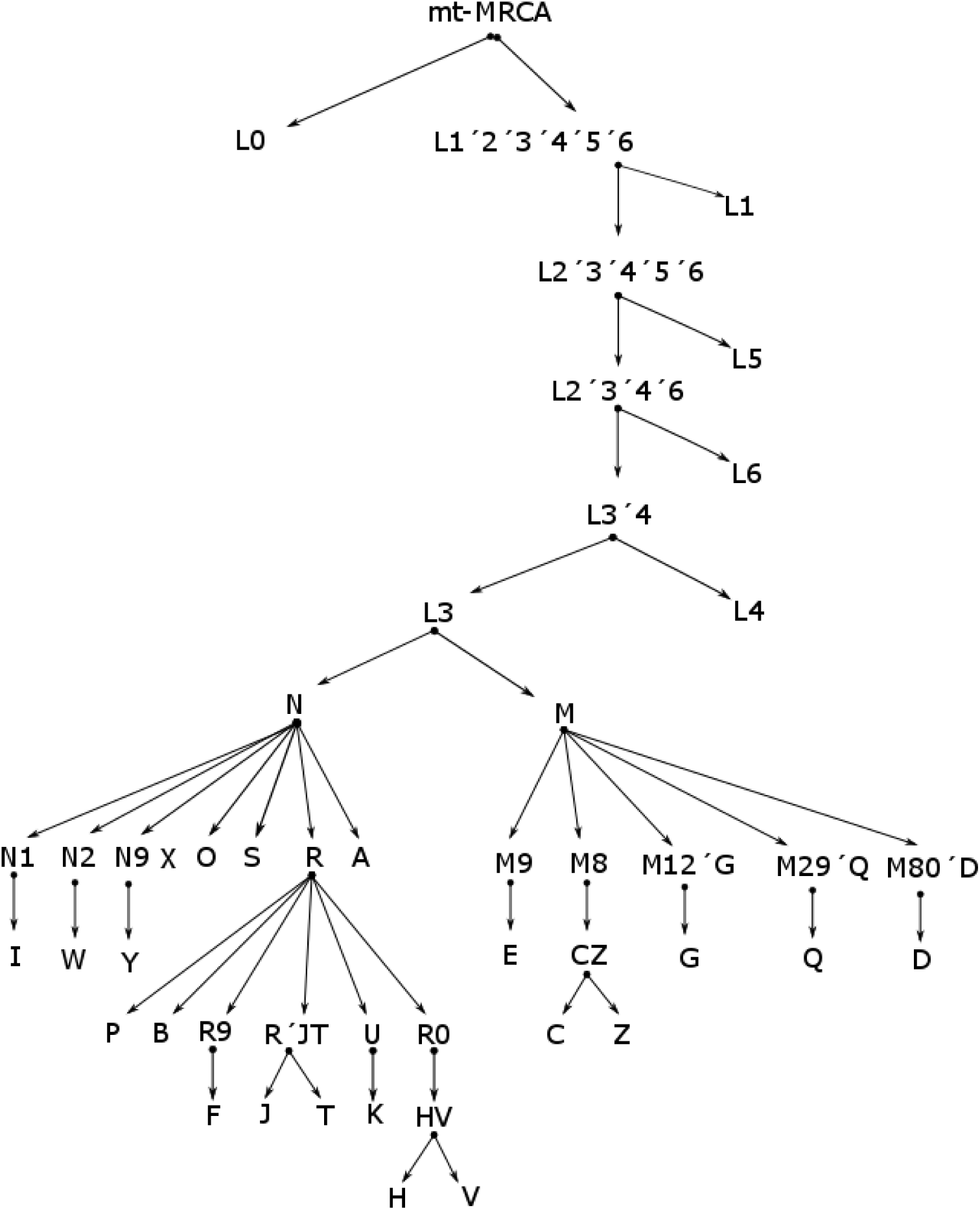
Phylogenetic tree of mtDNA sequences, modified from (www.phylotree.org). MRCA: Most recent common ancestor.

Mitochondrial hgs have been assigned a role as disease modifiers^5^. Particularly in neurological degenerative diseases such as Alzheimer’s disease ^20-23^ and Parkinson’s disease ^23-25^, but also in psychiatric disease^26^ and cardiac diseases such as hypertrophic^27^; ^28^ and ischemic cardiomyopathy^29^. Supporting a role as disease modifiers, some mtDNA hgs have specific physiological characteristics, e.g. reduced or increased ATP synthesis rates^30^; ^31^, and variation in methylation status of genes involved in inflammation and signaling^32^; ^33^. The association of mtDNA SNPs and hgs with both diseases and functional characteristics of mitochondria, has led to a pathogenic paradigm^34^ where variation in mitochondrial function is considered to be of paramount importance for development of disease. Specific hgs have also been associated with longevity^35^ and likelihood of being engaged in endurance athletic activities^36^. The clinical presentation of diseases caused by specific mtDNA variants depends, in some cases, on the hg background^37^. However, some of these studies are contradictory, either because they have been too poorly powered^38^, have not been carefully stratified with respect to sex, age, geographical background^39^ or population admixture^40^, or have used small areas of recruitment risking “occult” founder effects^41^. To circumvent some of these problems a recent large study on mtDNA SNPs identified a number of SNPs that were associated with several degenerative diseases^42^, however, the study pooled sequence information from a large geographical area, without correcting for potential population sub-structure.

Most countries have a complex history with repeated migrations^43^ and several bottle-necks caused by disease, war and emigration^44^; ^45^. These demographic events are reflected in the fine scale genetic structure within sub-populations^46^. However, the significance of such events to countrywide mtDNA hg distribution has not yet been assessed. As mtDNA interact functionally with the nuclear genome, it is paramount to ensure that specific mtDNA hgs – which are a marker of matrilineal genetic origin – do not represent population sub-structure at the genomic level. In theory, mtDNA is inherited independently of the nuclear genome, but population admixture and geographic isolation may result in linkage disequilibrium between mtDNA and the nuclear genome. Such a linkage disequilibrium might interfere with genetic association analysis.

Here we demonstrate the complexity of the spatio-temporal dynamics and genogeographic affinity of mtDNA hgs in 24,216 Danes, which were sampled at birth during a 25-year period. This number represents 1.6 % of the population. The sampling material was dried blood spots (DBSs) obtained as part of the Danish Neonatal Screening Program^47^, the very nature of which makes sampling geographically un-biased. Array analysis was performed using the PsychChip (Illumina, CA, USA) typing 588,454 variants.

## Materials and Methods

### Ethics statement

This is a register-based cohort study solely using data from national health registries. The study was approved by the Scientific Ethics Committees of the Central Denmark Region (www.komite.rm.dk) (J.nr.: 1-10-72-287-12) and executed according to guidelines from the Danish Data Protection Agency (www.datatilsynet.dk) (J.nr.: 2012-41-0110). Passive consent was obtained, in accordance with Danish Law nr. 593 of June 14, 2011, para 10, on the scientific ethics administration of projects within health research. Permission to use the DBS samples stored in the Danish Neonatal Screening Biobank (DNSB) was granted by the steering committee of DNSB (SEP 2012/BNP).

### Persons

As part of the iPSYCH (www.iPSYCH.au.dk) recruitment protocol, 24,651 singletons (47.1 % female), born between May 1 1981 and Dec 31 2005 were selected at random from the Danish Central Person Registry. The singletons had to have been alive one year after birth, and to have a mother registered in the Danish Central Person Registry. Furthermore, it should be possible to extract DNA from the DBS. DBS cards were obtained from the Danish Neonatal Screening Biobank at Statens Serum Institut ^48^ and DNA was extracted and analysed as described below. At the time of analysis (2012), the mean age of females was 18.2 years (SD: 6.6 years) and for males 18.8 years (SD: 6.7 years). There was no bias in the geographical distribution of the birthplace of samples, Figure 2.

**Figure 2.**
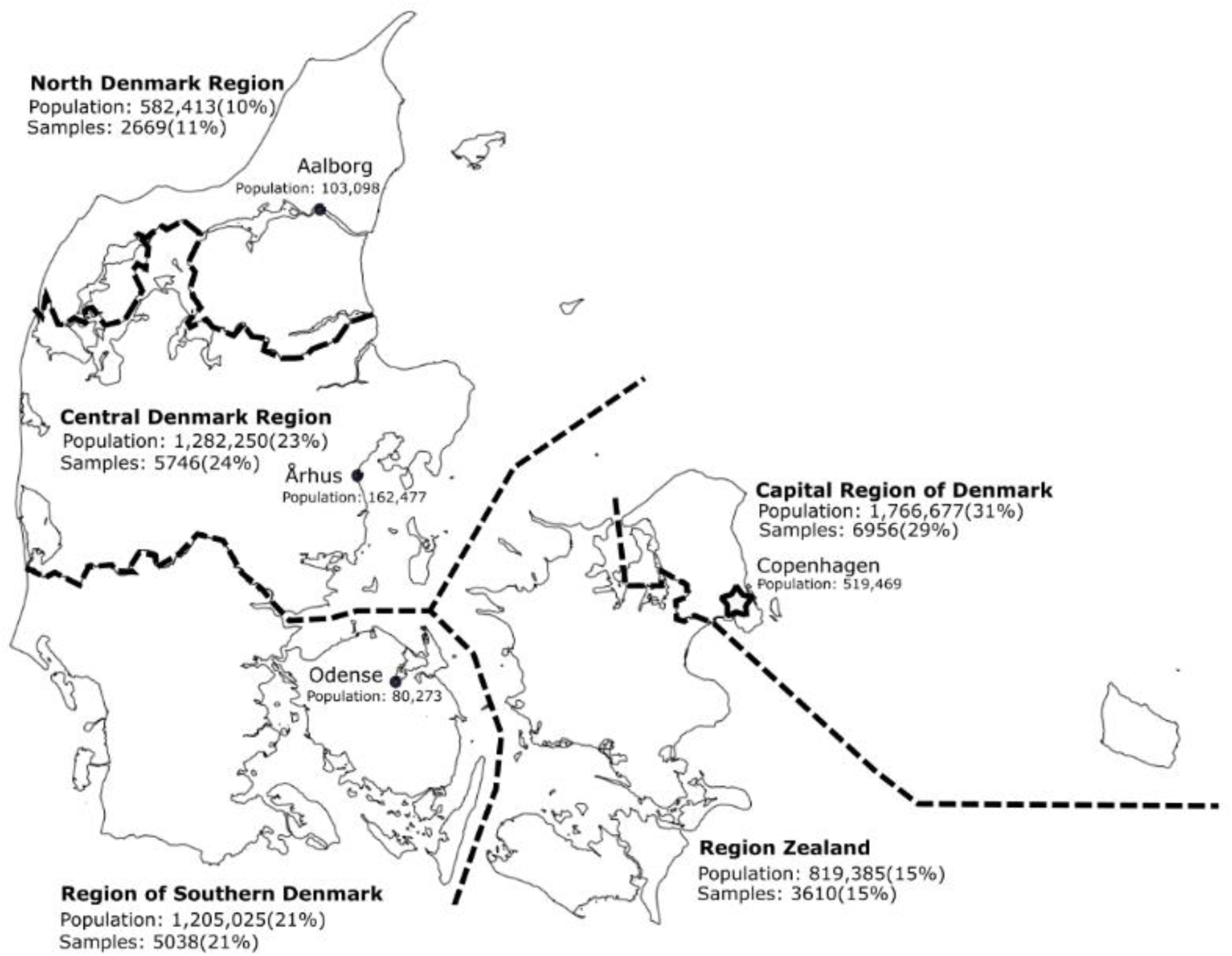
Map of Denmark with major administrative regions and metropolitan areas. The population of each region or area, as well as the number of samples, is given on the map. Based on birth-locations with ≥ 20 included individuals

### Genetic analysis

From each DBS card two 3.2-mm disks were excised from which DNA extracted using Extract-N-Amp Blood PCR Kit (Sigma-Aldrich, St Louis, MO, USA)(extraction volume: 200 μL). The extracted DNA samples were whole genome amplified (WGA) in triplicate using the REPLIg kit (Qiagen, Hilden, Germany), then pooled into a single aliquot. Finally, WGA DNA concentrations were estimated using the Quant-IT Picogreen dsDNA kit (Invitrogen, Carlsbad, CA, USA). The amplified samples were genotyped at the Broad Institute (MA, USA) using the Psychiatric Genetic Consortia developed PsychChip (Illumina, CA, USA) typing 588,454 variants. Following genotyping, samples with less than 97% call rate, as well as those where the estimated gender differed from the expected gender were removed from further analysis; altogether 435 (1.8 %) samples were removed due to problems with calling mtDNA variants. We then isolated the 418 mitochondrial loci and reviewed the genotype calls, before exporting into the PED/MAP format using GenomeStudio (Illumina, CA, USA). Samples were loaded into GenomeStudio (version 2011.a), a custom cluster was created using Gentrain (version 2), following automatic clustering all positions with heterozygotes were manually curated. The data was exported relative to the forward strand using PLINK Input Report Plug-in (version 2.1.3). Eigenvectors were calculated using PLINK (v1.90b3.31). PCA plots were created using the package ggplot2 (version 1.0.1) in R (version 3.1.3).

### mtDNA SNPing

Haplotyping of mtDNA was performed manually using the defining SNPs reported in www.phylotree.org ^19^. Hierarchical affiliation to macro-hg i.e. L0 – L6, M, N, R, and subsequent to hgs – units more distal in the cladogram, Figure 1, was performed. In some cases it was possible to establish affiliation to even sub-hgs. The call efficiencies of SNPs used in defining haplo- and sub-haplogroup affiliation are summarized in Suppl. Table 1.

### Phylogenetic analyses

Phylogenetic analyses was performed by constructing median-joining networks with Network 4.6.1.3 (http://www.fluxus-engineering.com). Fasta converted sequences of mtDNA SNPs from each person were aligned, sequences were pre-processed with Star Contraction (Maximum star radius 5 for R, N, M and 1 for L), then Median Joining networks were constructed (using the network parameters: Epsilon 10, Frequency >1, active) followed by post-processing with a maximum parsimony algorithm (MP) ^49^; ^50^. Network Publisher were used to post-process the networks^51^.

### Genogeographic affinity (nuclear genomic ancestry) analysis

Ancestry estimation was done using ADMIXTURE 1.3.0^52^. Briefly, a reference population consisting of Human Genome Diversity Project (HGDP) (http://www.hagsc.org/hgdp/) genotyping SNP data set, supplemented with representative samples of danes (716 individuals) and greenlanders (592 individuals) available at SSI from unrelated projects, was used. The final reference data set consisted of 103,268 SNPs and 2,248 individuals assigned to one of nine population groups: Africa, America, Central South Asia, Denmark, East Asia, non-Danish Europe, Greenland, Middle East and Oceania. K – number of clusters defined - was set to eight, based on principal component analysis clustering (data not shown).

Individuals belonging to individual mtDNA hgs or sub-hgs were merged with the reference population data set and analysed using ADMIXTURE. For prediction of the ancestry of individuals within the mtDNA hgs we created a random forest model^53^ based on the reference data set, with the clusters Q1-8 as predictors and population groups as outcome. The prediction was thus supervised. Prediction was done in R version 3.2.2, using the Caret package. The distribution of the eight basic clusters in samples of different geographical origin is shown in Suppl Figure 1. As expected, the African-characteristic cluster distribution plays a decreasing role when going from Africa over the Middle East to Central South Asia. Likewise, the Danish cluster distribution is very similar to that of Europe.

### Statistics

The statistical significance of differences in mtDNA proportions was assessed using a permutation version of Fisher’s exact test^54^. Calculations were performed using R ^55^. To assess population stratification^56^, principal component analysis (PCA) was performed.

## Results

### Distribution of macro-haplogroups

The mtDNA macro-hg distribution pattern is typically northern European^57^ with > 90% belonging to the R macro-hg, 7.0 % belonging to N and 1.6 % to M of likely Near Eastern or Asian origin^57^ and 0.7 % belonging to the combined L macro-hgs (L0-L6) of a likely African origin^58^, Table 1. A PCA analysis (PC 1 versus PC 2) based on all the SNPs showed a clear separation between R and L, with N and M located intermediately, figure 3A. The PC1 seems to reflect time since branching, whereas PC2 reflects geographical distance.

**Table 1.** Distribution of mtDNA macro hgs L0-L6, M, N and R. Incomplete is the number of individuals that could not be haplotyped using the haplotyping algorithm (see Materials and Methods).

| Macro-hg | n | % |
| --- | --- | --- |
| L0 | 26 | 0.1 |
| L1 | 13 | 0.1 |
| L2 | 54 | 0.2 |
| L3 | 71 | 0.3 |
| L4 | 16 | 0.1 |
| L5 | 2 | 0.0 |
| L6 | 0 | 0.0 |
| M | 394 | 1.6 |
| N | 1,700 | 7.0 |
| R | 21,940 | 90.6 |
| Total haplotyped | 24,216 | 100 |
| Incomplete | 426 | 1.7 |

### Distribution of haplogroups

The R macro-hg was dissolved into hgs as shown in Table 2. However, it was not possible, with the available SNPs, to differentiate the HV and P hgs from the R macro-hg (Suppl. Table 1, for details). Principle Component 1 versus 2, Figure 3B, demonstrated a clear clustering of mtDNA SNPs from persons belonging to each hg. The proximity of the clusters comprising U and K and H and V, respectively, is in accordance with the current phylogenetic mtDNA tree, Figure 1. Likewise, the median-joining graph of the R macro-hg, Figure 4A, based on all the called SNPs, disclosed a phylogenetic relationship compatible with that of Figure 1. In addition, all the hgs exhibit a considerable complexity, suggesting that each hg is the result of multiple migratory events and thus, multiple founder events. The sub-hg distribution of each of the hgs of the R macro-hg is shown in Table 3.

**Table 2.** Distribution of mtDNA haplogroups constituting the macro-hg R. Incomplete is the number of individuals that could not be haplotyped using the haplotyping algorithm (see Materials and Methods).

| **Hg** | **n** | **%** |
| --- | --- | --- |
| **H** | 10,912 | 50.0 |
| **U** | 3,237 | 14.8 |
| **T** | 2,323 | 10.6 |
| **J** | 2,195 | 10.1 |
| **K** | 1,754 | 8.0 |
| **V** | 785 | 3.6 |
| **R** | 570 | 2.6 |
| **F** | 50 | 0.2 |
| **B** | 10 | 0.0 |
| **Total haplotyped** | 21,836 | 100 |
| **Incomplete** | 104 | 0.5 |

**Figure 3.**
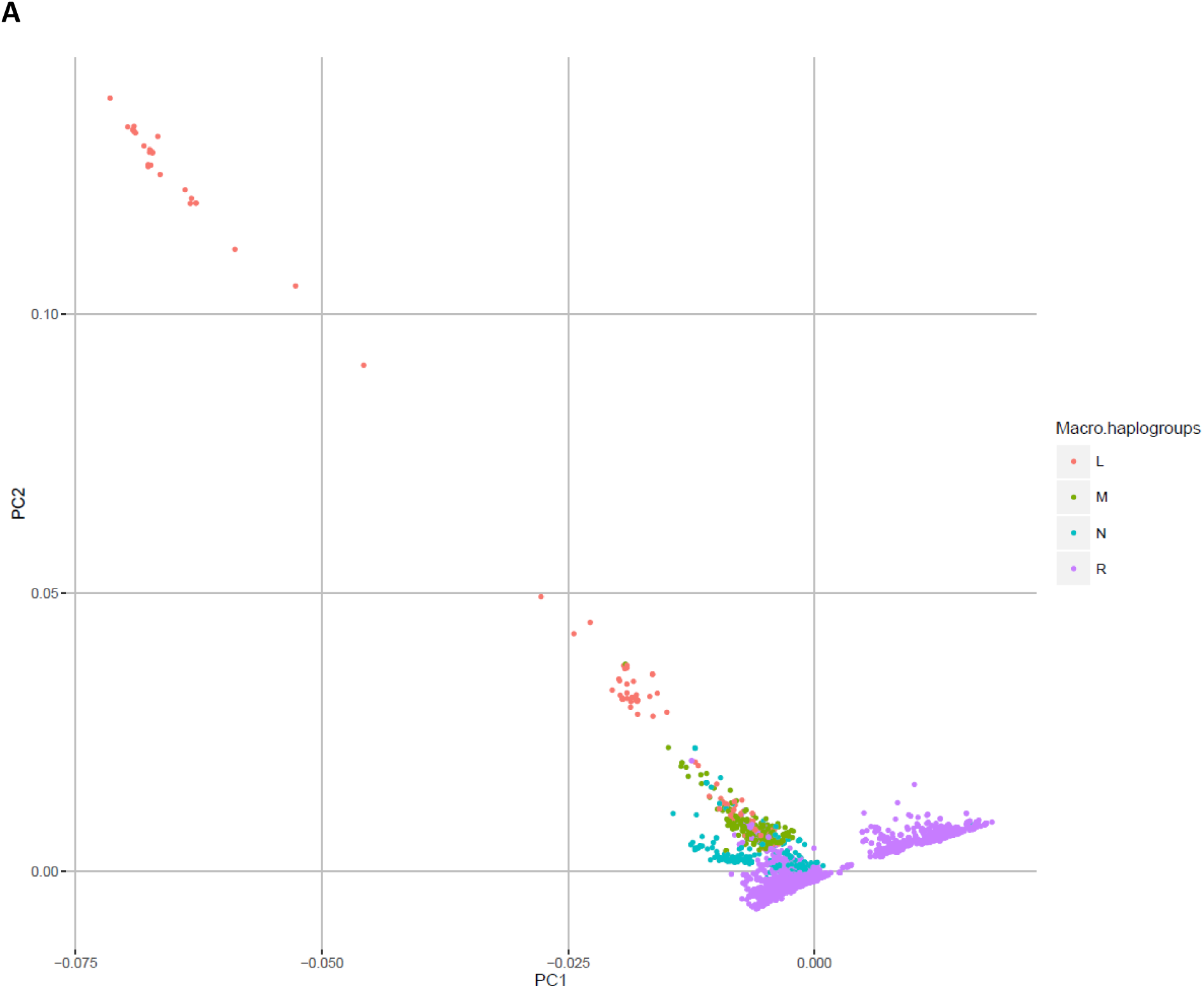
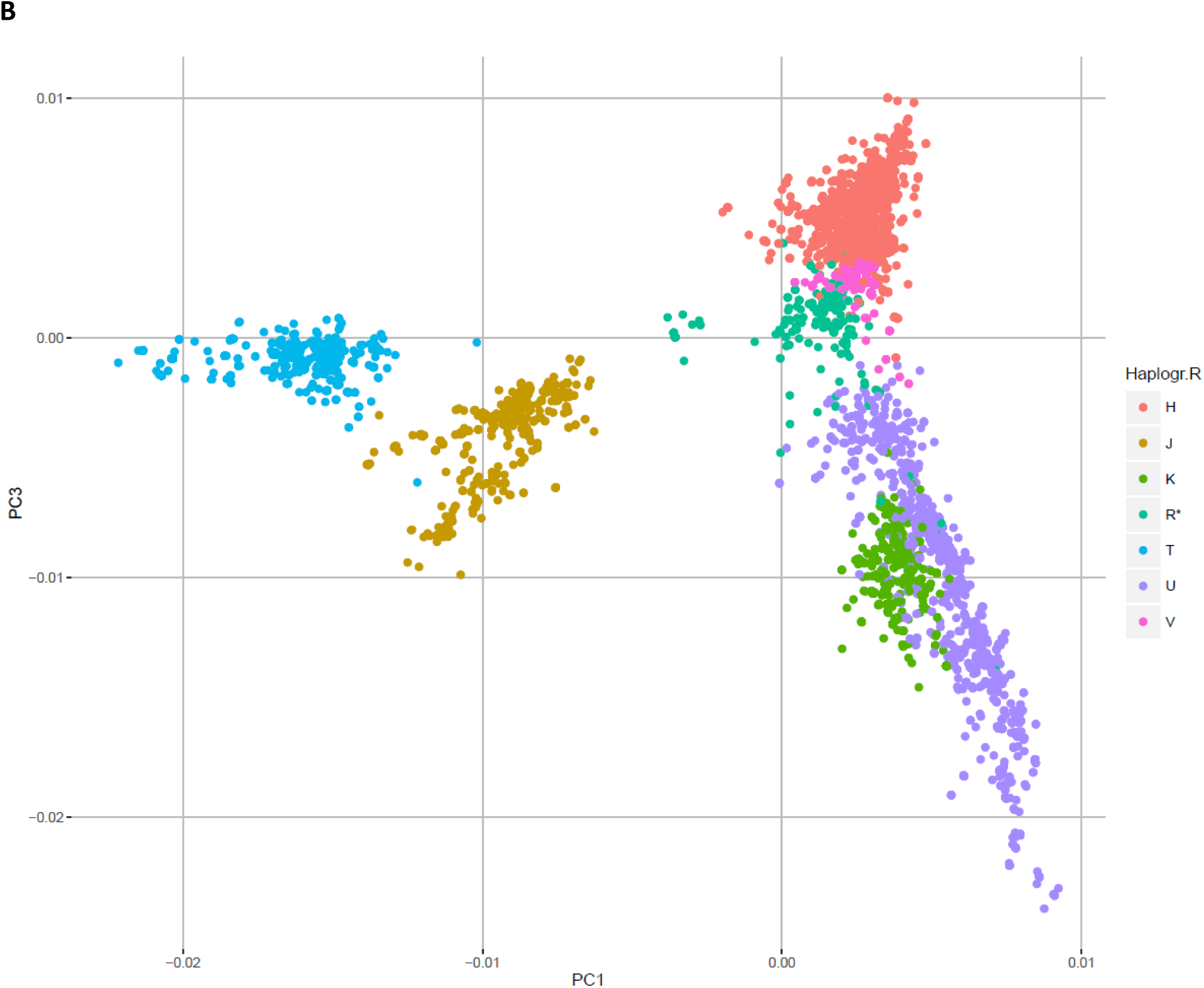
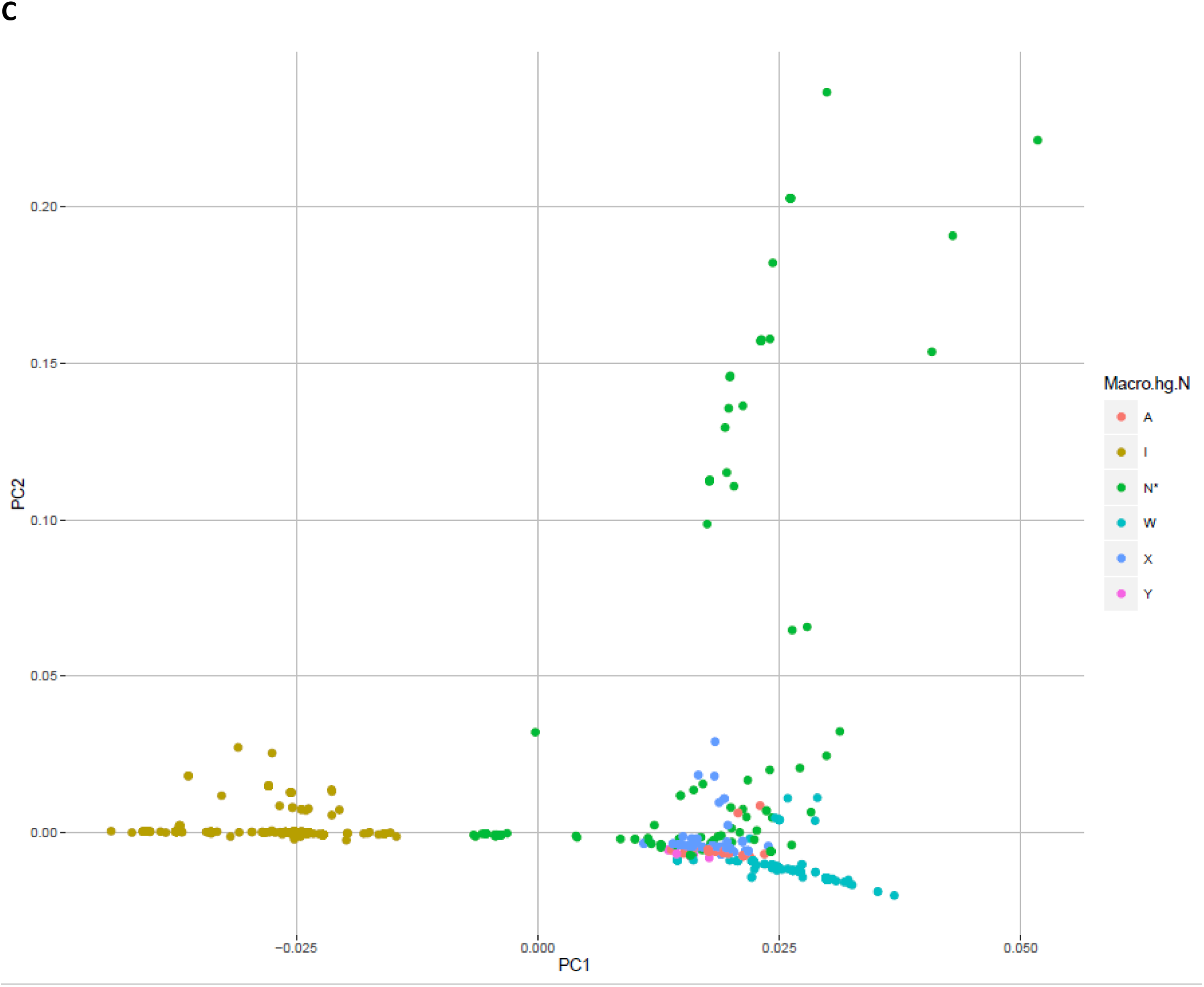
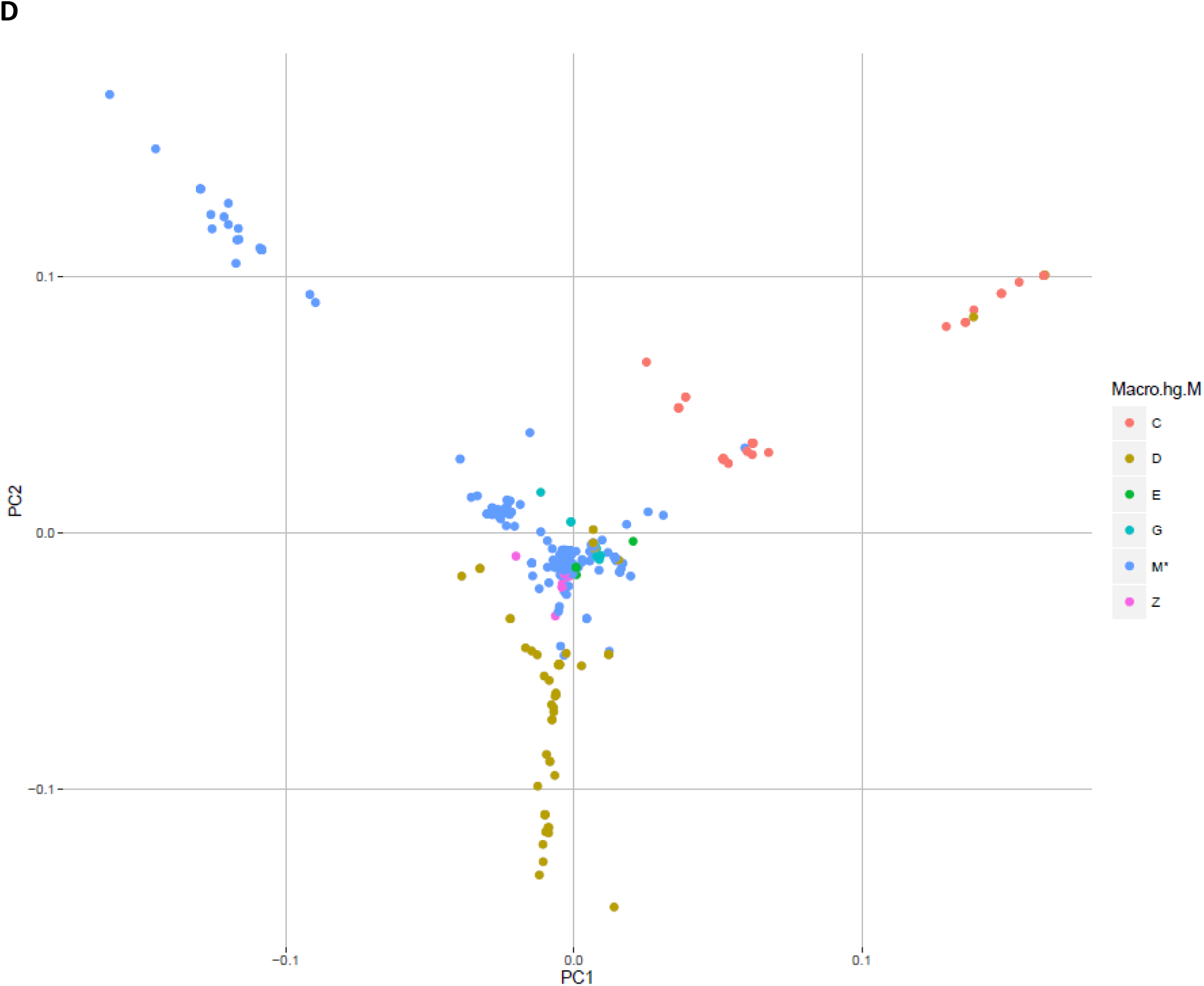
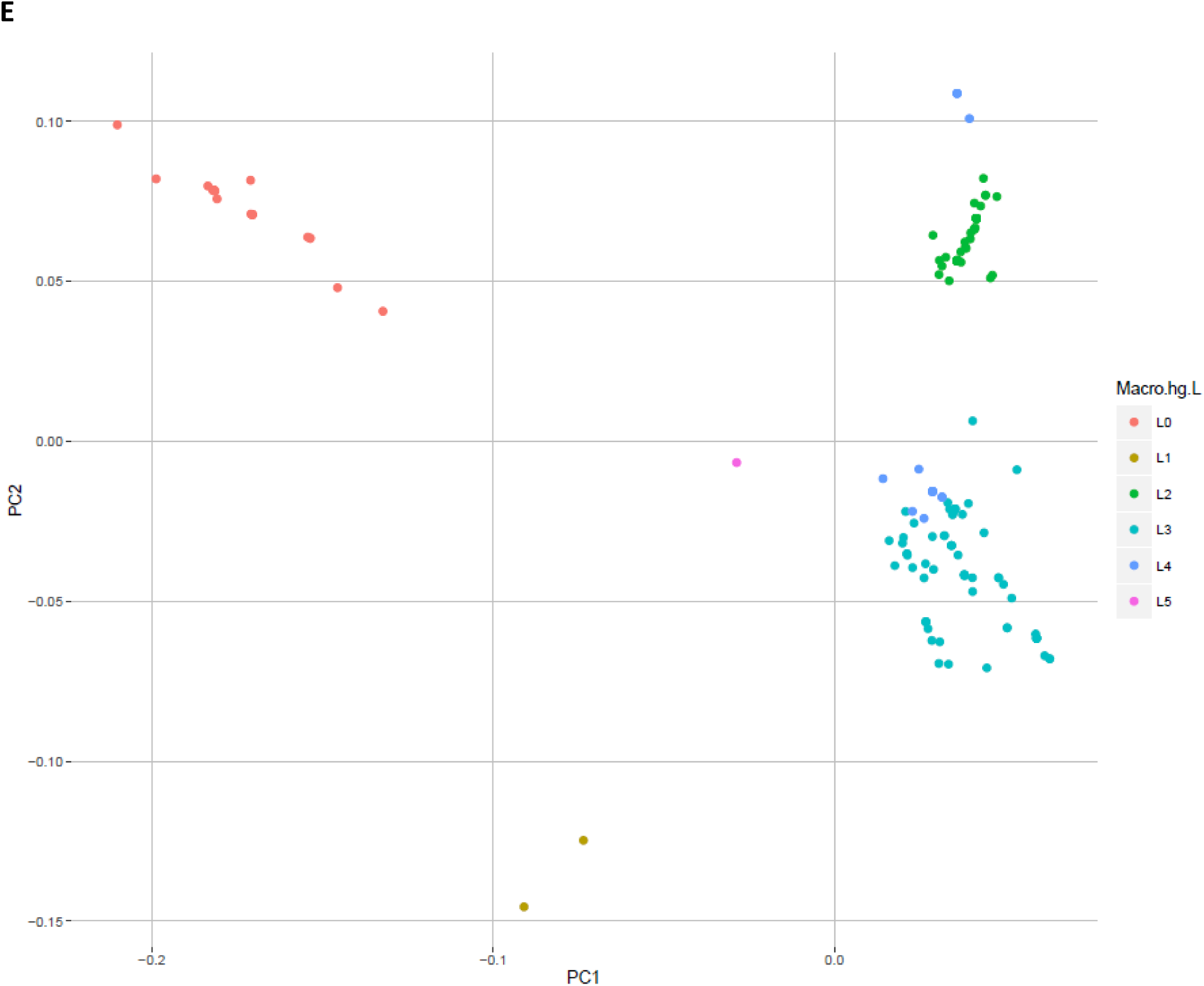
**A.** PCA of the macro-haplogroups L, M, N and R. **B.** PCA of the macro-haplogroup R, i.e. the major European haplogroups. **C**. PCA of the haplogroups belonging to macro-haplogroup N, and **D**. macro-haplogroup M, and **E**. PCA of the haplogroups belonging to macro-haplogroup L, i.e. the major African haplogroups. PC1: First principal component. PC2: Second principal component.

**Figure 4.**
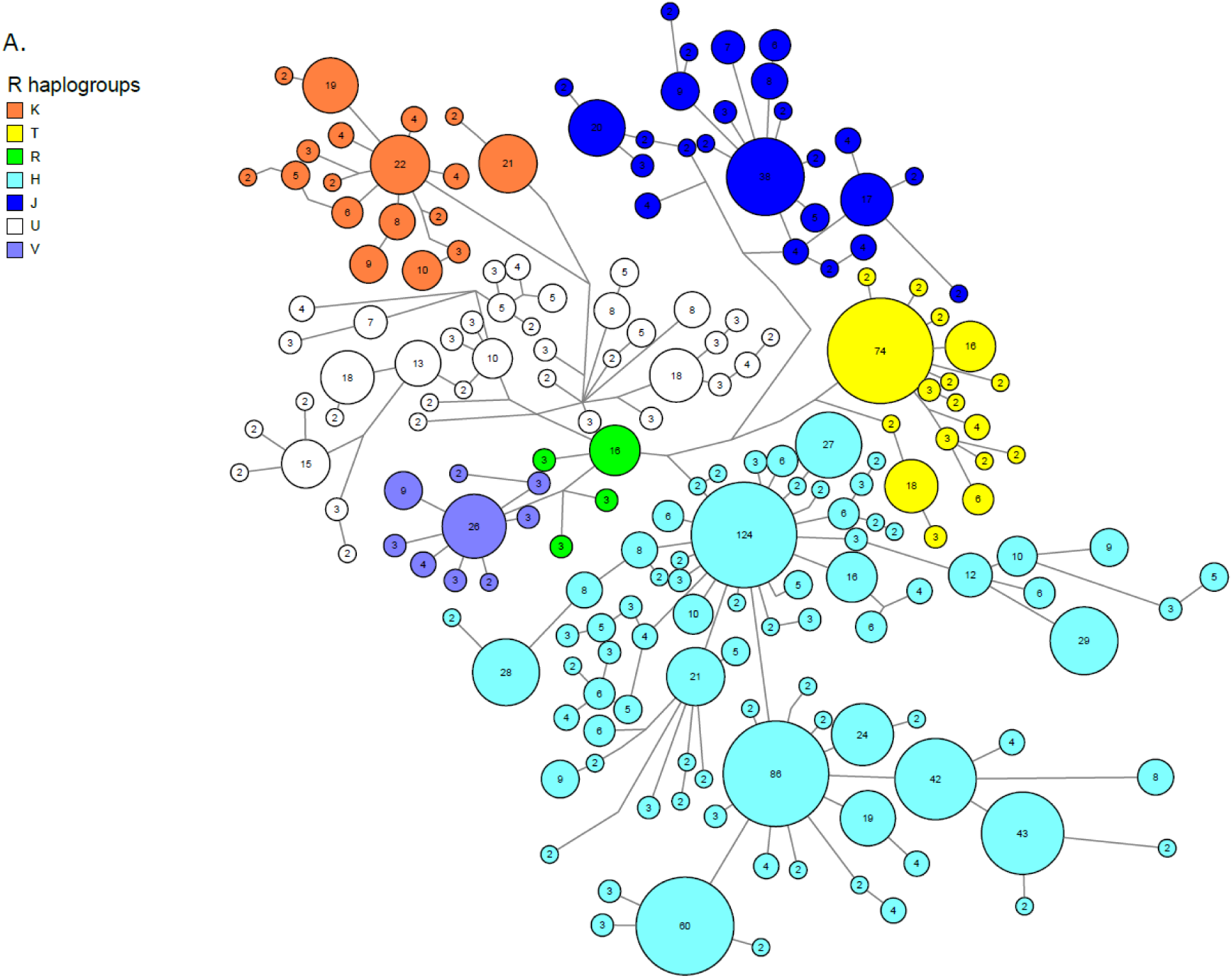
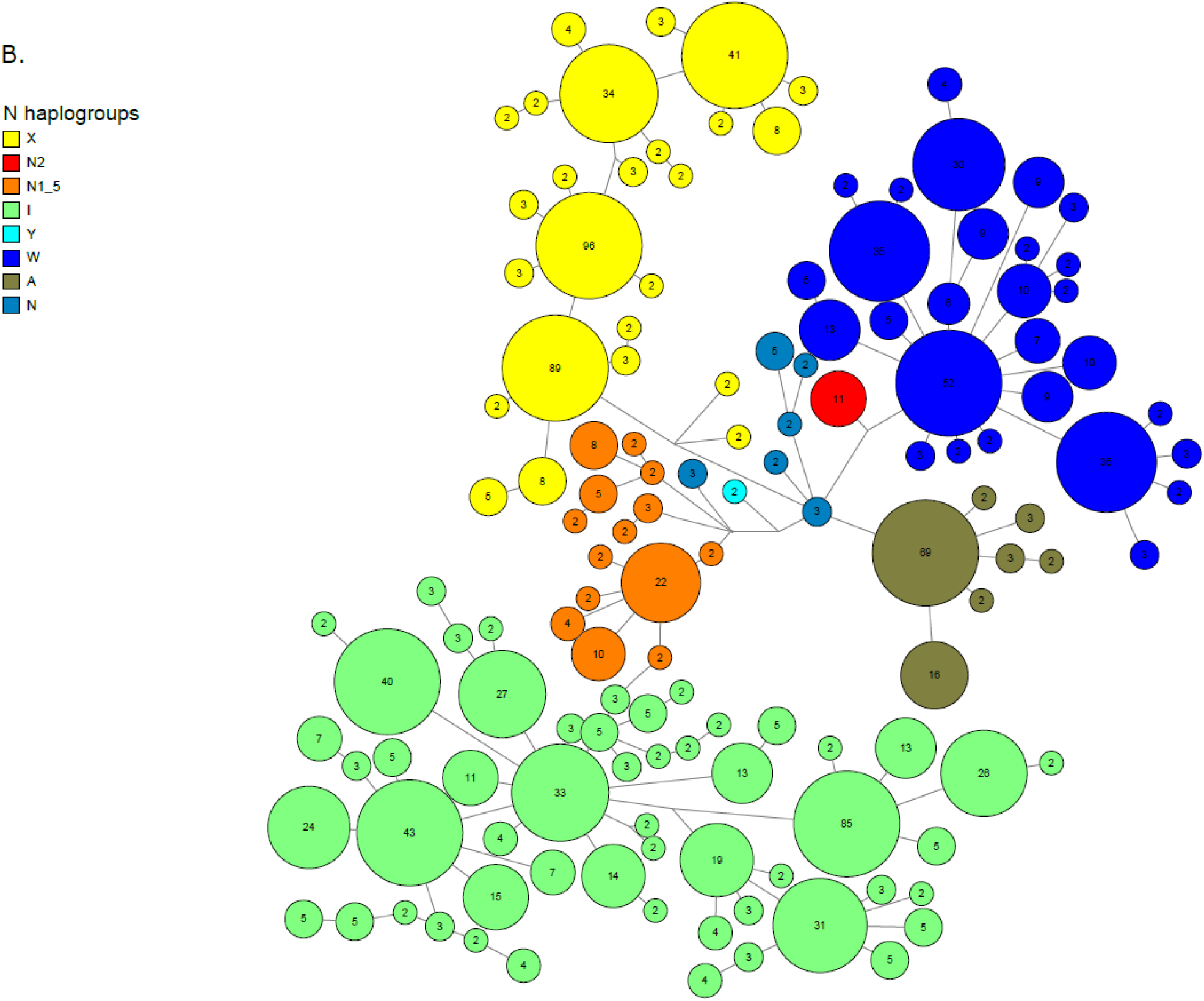
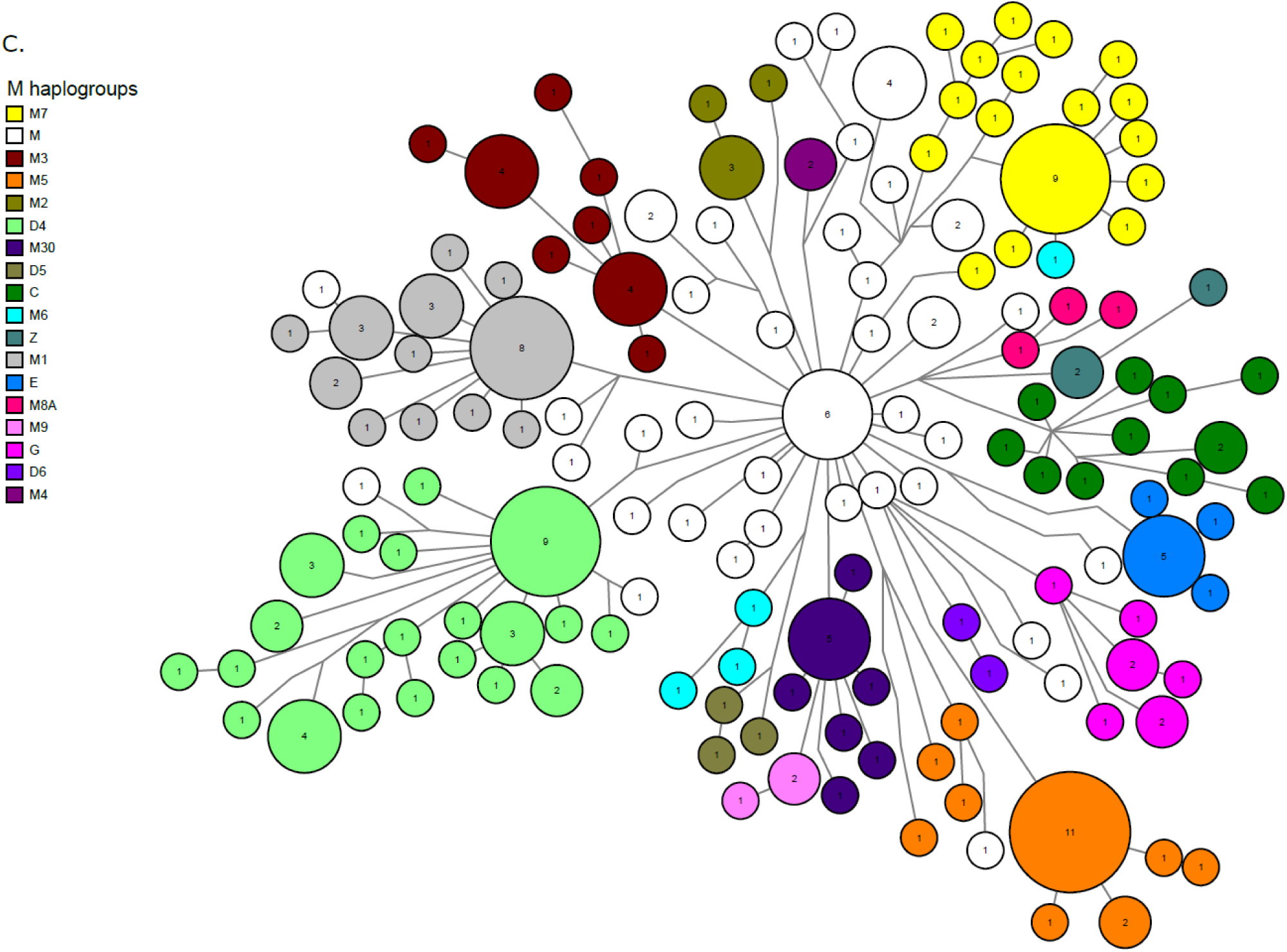
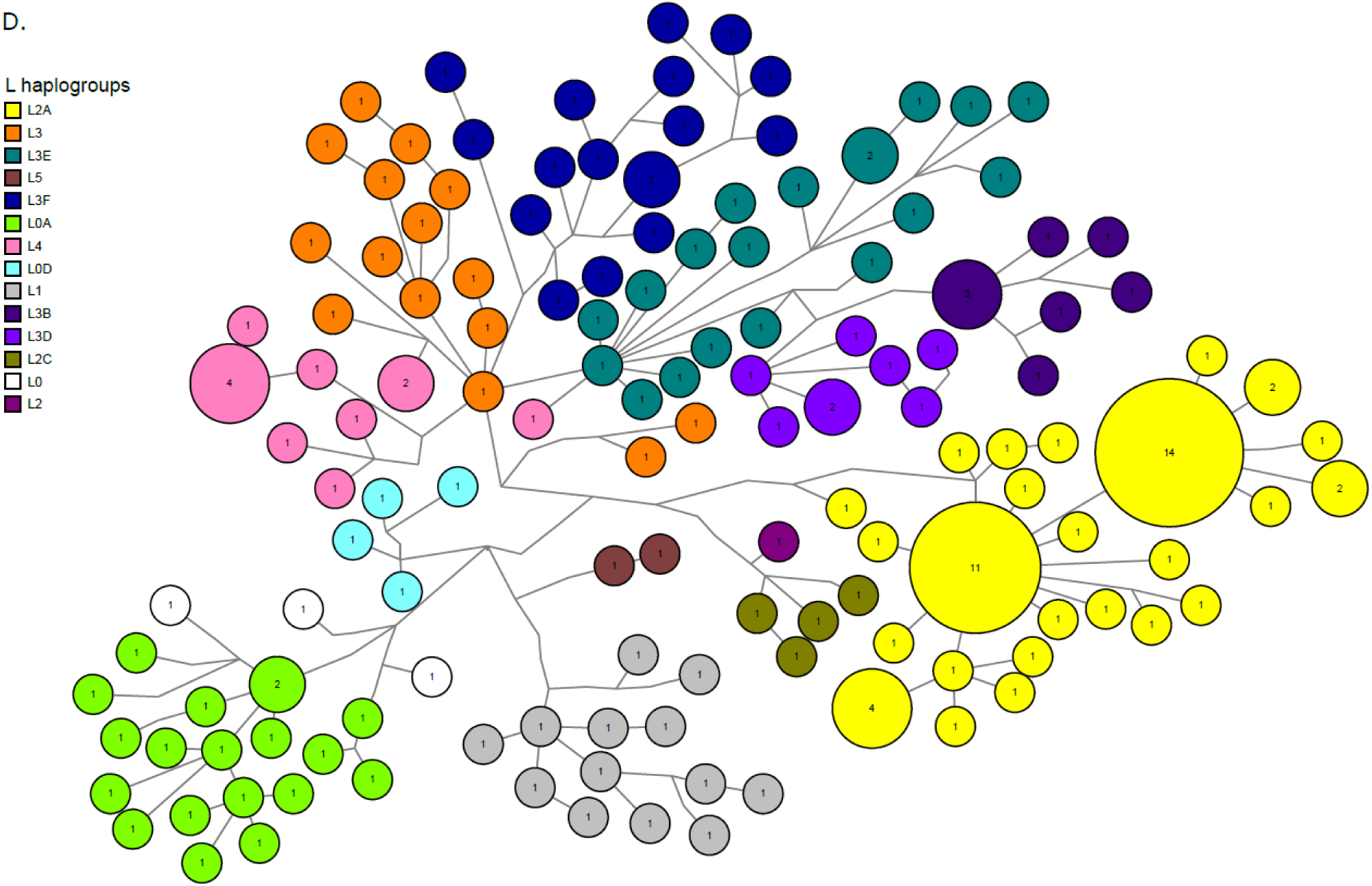
**A.** Median-joining (M-J) network of the macro-haplogroup R mtDNA sequences (2000 chosen at random). **B.** M-J network of the N and **C**. M haplogroups. **D**. M-J network of the L haplogroup sequences.

**Table 3.** Distribution of mtDNA sub-haplogroups of H, V, J, T, K, and U, the most frequent European haplogroups. HV could not be defined and is included in the H hg. Not sub-haplotyped is the number of individuals that could not be subhaplotyped using the algorithm defined by the SNPs in Suppl Table 1.

| Hg | Sub-hgs | n | % |
| --- | --- | --- | --- |
| **H** | H | 3,924 | 36.4 |
|  | H1-H30b-H79a | 4,054 | 37.6 |
|  | H2 | 1,058 | 9.8 |
|  | H3 | 699 | 6.5 |
|  | H4 | 220 | 2.0 |
|  | H5-36 | 828 | 7.7 |
|  | Number sub-haplotyped | 10,783 | 100 |
|  | Not sub-haplotyped | 129 | 1.2 |
| **U** | U | 120 | 3.8 |
|  | U1 | 66 | 2.1 |
|  | U2-U3 | 596 | 18.8 |
|  | U4-9 | 603 | 19.0 |
|  | U5a | 1,113 | 35.1 |
|  | U5b | 471 | 14.8 |
|  | U6 | 26 | 0.8 |
|  | U7 | 87 | 2.7 |
|  | U8 | 90 | 2.8 |
|  | Number sub-haplotyped | 3,172 | 100 |
|  | Not sub-haplotyped | 65 | 2.0 |
| **K** | K | 1 | 0.1 |
|  | K1 | 1,385 | 81.8 |
|  | K2 | 307 | 18.1 |
|  | K3 | 1 | 0.1 |
|  | Number sub-haplotyped | 1,694 | 100 |
|  | Not sub-haplotyped | 60 | 3.4 |
| **J** | J | 2 | 0.1 |
|  | J1 | 1,686 | 80.5 |
|  | J2 | 407 | 19.4 |
|  | Number sub-haplotyped | 2,095 | 100 |
|  | Not sub-haplotyped | 100 | 4.6 |
| **T** | T* | 9 | 0.4 |
|  | T1 | 396 | 17.1 |
|  | T2 | 1,908 | 82.5 |
|  | Number sub-haplotyped | 2,313 | 100 |
|  | Not sub-haplotyped | 10 | 0.4 |

The L, N and M macro-hgs were infrequent, Table 1. M and N could be broken down into hgs as shown in Table 4. The complexity of these macro-hgs, and macro-hg L, as demonstrated by median-joining phylogenetic analysis, Figure 4 B, C and D, suggest that these macro-hgs are composed of specific haplotypes from multiple immigrations over time. The number of DNA variations between individual haplotypes is so large that they can not represent development from a single founder within the time frame defined by the length of time in which Denmark has been populated.

**Table 4.** Distribution of mtDNA haplogroups contained within the N and M macro hgs. Not haplotyped is the number of individuals that could not be haplotyped using the algorithm defined by the SNPs in Suppl Table 1.

| **Macro-hg** | **(Sub) hgs** | **n** | **%** |
| --- | --- | --- | --- |
| **N** | A | 119 | 7.1 |
|  | I | 655 | 39.2 |
|  | N | 55 | 3.3 |
|  | N1-5 | 111 | 6.6 |
|  | N2 | 11 | 0.7 |
|  | W | 319 | 19.1 |
|  | X | 394 | 23.6 |
|  | Y | 6 | 0.4 |
|  | Number haplotyped | 1,670 | 100 |
|  | Not haplotyped | 30 | 1.8 |
| **M** | C | 40 | 10.7 |
|  | D4 | 53 | 14.2 |
|  | D5 | 4 | 1.1 |
|  | D6 | 2 | 0.5 |
|  | E | 8 | 2.1 |
|  | G | 10 | 2.7 |
|  | M | 81 | 21.7 |
|  | M1 | 30 | 8.0 |
|  | M2 | 11 | 2.9 |
|  | M3 | 24 | 6.4 |
|  | M30 | 14 | 3.7 |
|  | M4 | 3 | 0.8 |
|  | M5 | 26 | 7.0 |
|  | M6 | 9 | 2.4 |
|  | M7 | 37 | 9.9 |
|  | M8a | 7 | 1.9 |
|  | M9 | 4 | 1.1 |
|  | Z | 11 | 2.9 |
|  | Number haplotyped | 374 | 100 |
|  | Not haplotyped | 20 | 5,1 |

### Geno-geographical affinity of mtDNA haplogroups

An admixture analysis of the persons of different hgs was performed with results as shown in Table 5. The major hgs, H and its sub-hgs, have a 90-95% Danish ancestry and ~ 5% non-Danish European structure. However, there is a great variation between mtDNA hgs – most pronounced in hg U -, and most of the hgs have a notable – but varying – proportion of admixture from Europe, Middle East and Central South Asia, Table 5. This finding is compatible with the very complex M-J diagram of the Danish hgs, Figure 4A. The ancestry or geno-geographic affinity of the macro-hgs M and N differ even more, Table 5, with the M being of South East and East Asian affinity, and N of Danish and European affinity. Macro-hg L exhibited – surprisingly – a predominance of Middle Eastern and European genomic affinity, Table 5, where an African ancestry should be expected^57^; ^59^.

**Table 5.**
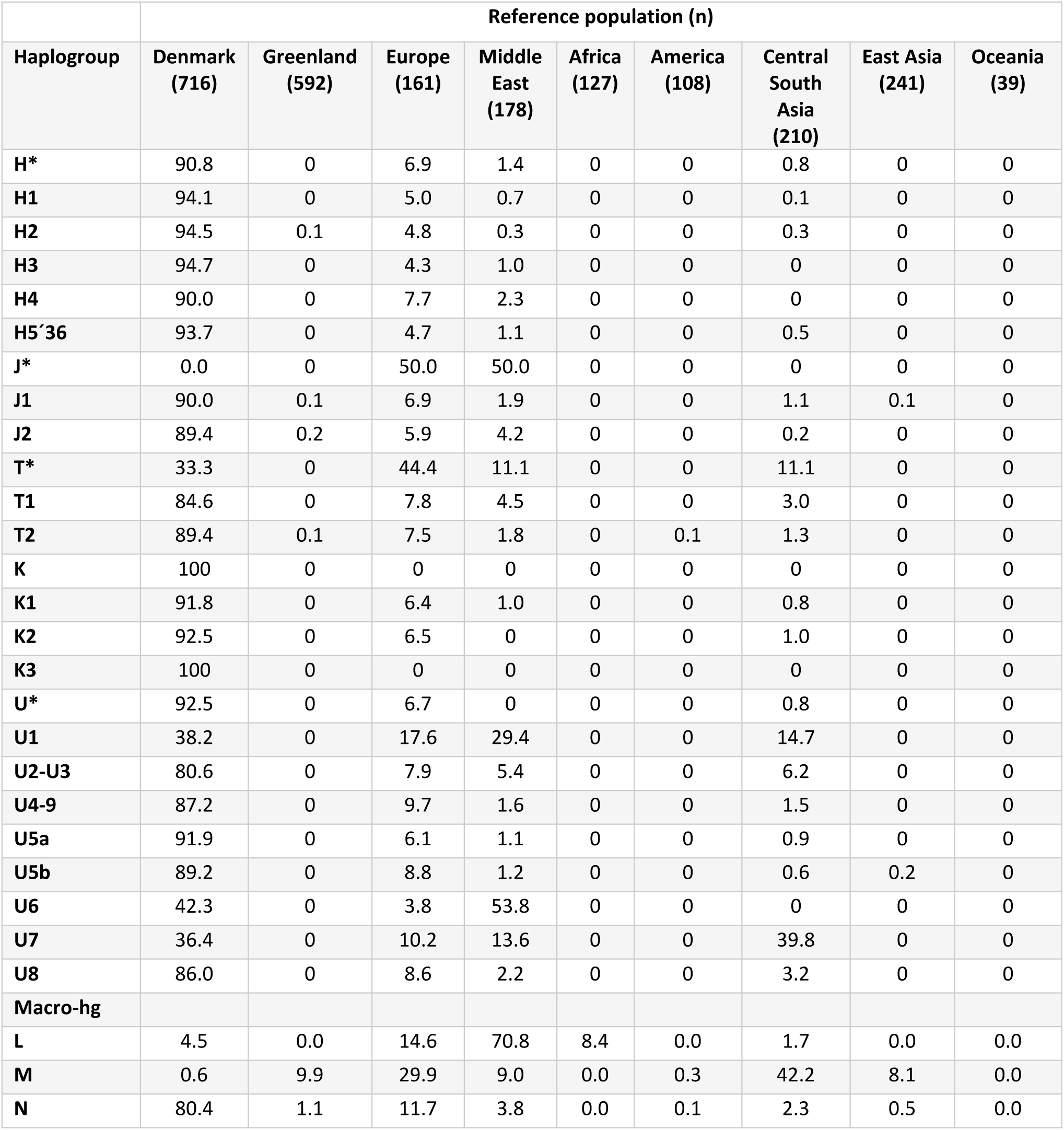
Geno-geographic affinity (ancestry) of the autosomal genome of persons in the study broken down with respect to mtDNA macro-hg or hg. The number of persons in each geographical group in the reference population is given in the header.

| Haplogroup | Reference population (n) |  |  |  |  |  |  |  |  |
| --- | --- | --- | --- | --- | --- | --- | --- | --- | --- |
|  | Denmark<br>(716) | Greenland<br>(592) | Europe<br>(161) | Middle<br>East<br>(178) | Africa<br>(127) | America<br>(108) | Central<br>South<br>Asia<br>(210) | East Asia<br>(241) | Oceania<br>(39) |
| **H\*** | 90.8 | 0 | 6.9 | 1.4 | 0 | 0 | 0.8 | 0 | 0 |
| **H1** | 94.1 | 0 | 5.0 | 0.7 | 0 | 0 | 0.1 | 0 | 0 |
| **H2** | 94.5 | 0.1 | 4.8 | 0.3 | 0 | 0 | 0.3 | 0 | 0 |
| **H3** | 94.7 | 0 | 4.3 | 1.0 | 0 | 0 | 0 | 0 | 0 |
| **H4** | 90.0 | 0 | 7.7 | 2.3 | 0 | 0 | 0 | 0 | 0 |
| **H5'36** | 93.7 | 0 | 4.7 | 1.1 | 0 | 0 | 0.5 | 0 | 0 |
| **J\*** | 0.0 | 0 | 50.0 | 50.0 | 0 | 0 | 0 | 0 | 0 |
| **J1** | 90.0 | 0.1 | 6.9 | 1.9 | 0 | 0 | 1.1 | 0.1 | 0 |
| **J2** | 89.4 | 0.2 | 5.9 | 4.2 | 0 | 0 | 0.2 | 0 | 0 |
| **T\*** | 33.3 | 0 | 44.4 | 11.1 | 0 | 0 | 11.1 | 0 | 0 |
| **T1** | 84.6 | 0 | 7.8 | 4.5 | 0 | 0 | 3.0 | 0 | 0 |
| **T2** | 89.4 | 0.1 | 7.5 | 1.8 | 0 | 0.1 | 1.3 | 0 | 0 |
| **K** | 100 | 0 | 0 | 0 | 0 | 0 | 0 | 0 | 0 |
| **K1** | 91.8 | 0 | 6.4 | 1.0 | 0 | 0 | 0.8 | 0 | 0 |
| **K2** | 92.5 | 0 | 6.5 | 0 | 0 | 0 | 1.0 | 0 | 0 |
| **K3** | 100 | 0 | 0 | 0 | 0 | 0 | 0 | 0 | 0 |
| **U\*** | 92.5 | 0 | 6.7 | 0 | 0 | 0 | 0.8 | 0 | 0 |
| **U1** | 38.2 | 0 | 17.6 | 29.4 | 0 | 0 | 14.7 | 0 | 0 |
| **U2-U3** | 80.6 | 0 | 7.9 | 5.4 | 0 | 0 | 6.2 | 0 | 0 |
| **U4-9** | 87.2 | 0 | 9.7 | 1.6 | 0 | 0 | 1.5 | 0 | 0 |
| **U5a** | 91.9 | 0 | 6.1 | 1.1 | 0 | 0 | 0.9 | 0 | 0 |
| **U5b** | 89.2 | 0 | 8.8 | 1.2 | 0 | 0 | 0.6 | 0.2 | 0 |
| **U6** | 42.3 | 0 | 3.8 | 53.8 | 0 | 0 | 0 | 0 | 0 |
| **U7** | 36.4 | 0 | 10.2 | 13.6 | 0 | 0 | 39.8 | 0 | 0 |
| **U8** | 86.0 | 0 | 8.6 | 2.2 | 0 | 0 | 3.2 | 0 | 0 |
| **Macro-hg** |  |  |  |  |  |  |  |  |  |
| **L** | 4.5 | 0.0 | 14.6 | 70.8 | 8.4 | 0.0 | 1.7 | 0.0 | 0.0 |
| **M** | 0.6 | 9.9 | 29.9 | 9.0 | 0.0 | 0.3 | 42.2 | 8.1 | 0.0 |
| **N** | 80.4 | 1.1 | 11.7 | 3.8 | 0.0 | 0.1 | 2.3 | 0.5 | 0.0 |

### Spatial distribution of mtDNA haplogroups

Denmark is divided into five geographical and administrative regions as shown in Figure 2. The samples for this study were obtained from all regions in Denmark and linked to the postal code of the birthplace. The frequency of the macro-hgs and the most frequent hgs is shown for each administrative region in Table 6. The most marked differences were the relatively low frequency of hg H, 43.4 %, in the Capital Region, compared to the North Denmark Region with 48.8 % hg H (p < 0.0001), and the higher frequencies of hgs L, M and R (in total 7.5 %) in the Capital Region compared to the North Denmark Region (in total 3.9 %) (p < 0.0001). Thus, in the Capital Region, the L hgs have a frequency of 1.3 %, compared with 0.4 – 0.6 % in the other regions and the M hgs have a frequency of 2.6 % in the Capital region compared to 1.0 – 1.6 % in the other regions. As the L and M hgs are rare in the European population and very frequent in African and Asian populations, the noted difference probably reflects a higher proportion and preferential localization of non-ethnic Danes in the Capital region. A similar spatial difference was apparent when the hg distributions of persons from the Danish metropolitan areas, comprising the major cities, Copenhagen, Aarhus, Odense and Aalborg, were compared with the distributions in the remaining rural areas, Suppl Table 2. In the metropolitan areas, the combined frequency of L and M hhgs was 4.4 % as compared to 1.9 % in the rural areas.

**Table 6.** Distribution of the most frequent haplogroups in the five administrative regions of Denmark. Only persons born at locations with ≥ 20 births are included.

| **Haplogroup** | **Capital Region** | **Region Zealand** | **Central Denmark Region** | **Region of South Denmark** | **North Denmark Region** |
| --- | --- | --- | --- | --- | --- |
|  | % (n) | % (n) | % (n) | % (n) | %(n) |
| **H** | 43.4 (2,962) | 46.9 (1,664) | 45.7 (2,584) | 45.6 (2,259) | 48.8 (1,284) |
| **I** | 2.6 (175) | 2.5 (89) | 3.2 (180) | 3.0 (148) | 2.3 (60) |
| **J** | 8.6 (586) | 8.8 (313) | 9.4 (534) | 9.4 (464) | 8.2 (215) |
| **K** | 6.9 (468) | 6.3 (224) | 8.1 (456) | 7.3 (364) | 8.7 (229) |
| **L** | 1.3 (88) | 0.5 (17) | 0.5 (31) | 0.4 (20) | 0.6 (17) |
| **M** | 2.6 (176) | 1.3 (45) | 1.0 (56) | 1.6 (79) | 1.0 (27) |
| **N\*** | 1.3 (92) | 1.2 (43) | 1.2 (68) | 1.2 (61) | 1.3 (33) |
| **R** | 3.6 (244) | 2.1 (74) | 2.5 (144) | 2.4 (119) | 2.3 (61) |
| **T** | 9.2 (627) | 10.0 (354) | 9.3 (525) | 8.8 (434) | 9.0 (238) |
| **U** | 13.8 (939) | 13.8 (491) | 13.1 (744) | 13.4 (666) | 13.1 (346) |
| **V** | 3.4 (229) | 3.3 (116) | 3.4 (195) | 3.5 (175) | 2.8 (73) |
| **W** | 1.4 (96) | 1.6 (57) | 1.3 (74) | 1.4 (67) | 0.9 (24) |
| **X** | 2.0 (134) | 1.7 (60) | 1.2 (67) | 2.0 (101) | 0.9 (24) |
| **NA** | 0 (2) | 0.1 (3) | 0 (2) | 0 (1) | 0 (1) |
| **Total:** | 100 (6,818) | 100 (3,550) | 100 (5,660) | 100 (4,957) | 100 (2,632) |

### Temporal distribution of mtDNA haplogroups

The frequency of the major hgs H, J, T, K, V did not change significantly from year to year in the period from 1981 to 2005 (Results not shown).

The frequencies of M and L hgs, Figure 5, increased significantly over the period. The L hgs increased from a constant level of ~ 0.4 % from 1981 – 1995 to ~ 1.5 % in 2005. The M hgs rose from a constant level of ~ 1% from 1981 – 1991 to ~ 3 % in 2005. The proportion of L and M hgs increased significantly from the period 1981-1986 to the period 2000-2005, Table 7. The considerable diversity of the M macro-hg, Figure 4C and the extreme diversity of the L macro-hg, Figure 4D, as disclosed by the M-J-network, suggests that the hgs are the result of immigration from different source populations. The occurrence of new clusters on the PCA when the macro-hgs M and L from the period 1981-1986 were compared with the same macro-hgs in 2000-2005, Suppl. Figs. 2A & B, makes it likely that the increase in the proportion of both macro-hgs represent immigration.

**Figure 5.**
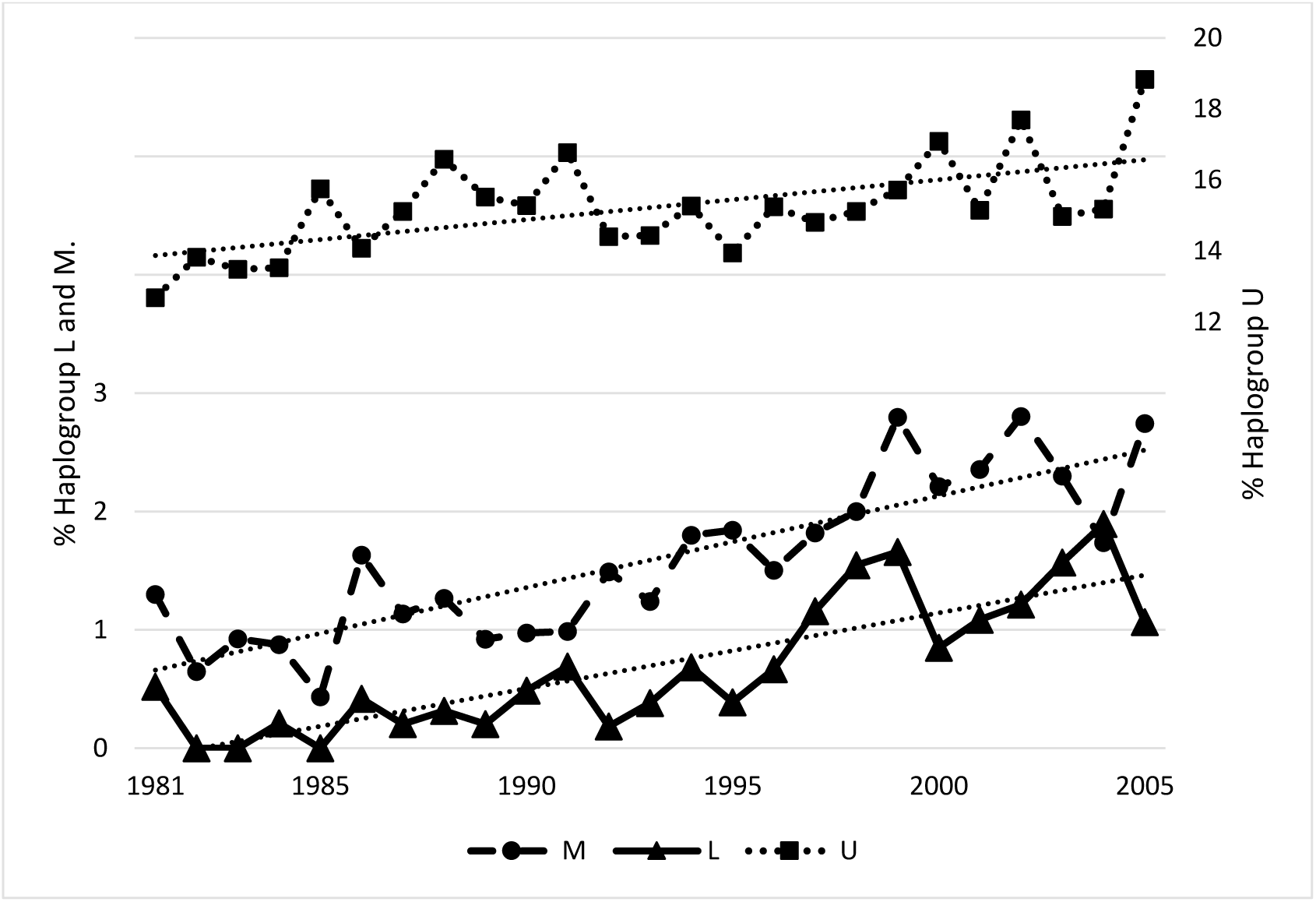
The proportion of haplogroup L, M and U as a function of birth year.

**Table 7.** Distributions of the most frequent haplogroups in the period 1981-1986 and 2000-2005.

| **Haplogroup** | **1981-1986** | **2000-2005** | **P-Value** |
| --- | --- | --- | --- |
|  | **% (n)** | **% (n)** |  |
| **H** | 47.3 (2,162) | 44.3 (2,454) | 2,98E-03 |
| **I** | 2.8 (130) | 2.7 (150) | 0,71 |
| **J** | 8.9 (405) | 9.2 (511) | 0,53 |
| **K** | 7.9 (359) | 7.0 (387) | 0,10 |
| **L** | 0.2 (8) | 1.2 (67) | 1,14E-10 |
| **M** | 1.0 (44) | 2.4 (131) | 3,73E-08 |
| **N\*** | 1.1 (50) | 1.3 (73) | 0,32 |
| **R** | 2.2 (102) | 2.9 (163) | 0,03 |
| **T** | 9.8 (447) | 8.5 (468) | 0,02 |
| **U** | 12.7 (579) | 14.3 (790) | 0,02 |
| **V** | 3.6 (163) | 3.1 (169) | 0,16 |
| **W** | 1.2 (56) | 1.4 (80) | 0,39 |
| **X** | 1.4 (64) | 1.6 (89) | 0,41 |
| **NA** | 0.0 (1) | 0.1 (3) | 0,63 |
| **Total** | 100 (4,570) | 100 (5,535) | - |

The hg U also increased in proportion each year from 1981 to 2005, Figure 5, albeit not significantly. However, when analysing only the R-macro-hg, the proportion of hg-U increased significantly (data not shown). In the case of the U-hg, the PCA did not reveal the appearance of novel clusters when comparing 1981-1986 with 2000-2005, Suppl Fig 2C. Thus, there was no evidence for the introduction of novel U-hgs over the period. However, a more detailed analysis revealed that the increase in U-hgs was largely due to an increase in the infrequent sub-hgs U1, U6, U7, and U8, Suppl Table 3, as all these increased by more than 40%. An admixture analysis showed that whereas the Danish autosomal genomic affinity of the hgs U*, U2-U3, U4-9, U5a&b is in the range of 80.6% - 92.5 %, Table 5, comparable to that of the other major European hgs, the Danish affinity drops to 36.4% - 42.3 % for U1, U6 and U7. According to Table 5, the U1 and U7 have strong Middle Eastern (29.4% and 13.6%) and Central South Asian (14.7% and 39.8%) autosomal geno-geographic affinities, and the U6 exhibits a very strong Middle Eastern autosomal affinity (53.8%). This suggests that the increase in frequency of hg U is largely due to expansion of hgs brought into Denmark as a result of recent immigration.

## Discussion

This study shows that the distribution of mtDNA hgs in Denmark is highly dynamic and complex. It comprises 1.6 % of the Danish population over a 25-year period, and is by far the largest performed of the distribution of mtDNA hgs in any country. The method of collecting stored DBS from the PKU biobank, where the coverage is ~ 99 % ^60^, enabled us to survey a true population based sample of mtDNA from persons, where the time and place of birth was known from national electronic registries. This is in contrast to most other population genetic studies of adults sampled in a specific bias-prone context, e.g. hospitalized patients or geographically biased samplings.

The distribution of macro-hgs, Table 1, is typical of Northern Europe^61^; ^62^. The large majority of persons had hgs belonging to the R macro-hg, Table 2. The distribution of hgs within the R macro-hg, Table 2, is very similar to that previously described among 9000+ persons from the greater Copenhagen area^63^ and a much smaller (~200 cases) Danish forensic control sample^64^, as well as that found in 2000 Danish exomes from a mixed control and diabetes population^65^. However, the latter study is seriously biased as it contains ~ 50% patients with type-2 diabetes, which have shown associations with mtDNA haplotypes^65^. Despite this, the sub-hg distribution in macro-hg R is similar to that seen in the present study, Table 3, e.g. H1 and H3 constitute 37.6 % and 6.5 % of the H hgs in our study and 39.6 % and 5.2 % in the Li et al. study^65^.

The mtDNA haplotyping was based on array data with only 418 mtDNA SNPs and as a consequence not all sub-hgs could be called. Ideally sub-haplotyping could be refined – without including sub-hg defining SNPs from Phylotree – by the application of clustering approaches. However, this was not attempted, as it would lead to results that could not be compared with other studies. The stringent adherence to specific SNPs meant that a number of persons, normally in the order of 1-2% (see tables 2-4), were not assigned to a specific haplotype. An advantage of using a limited set of markers is that confounding due to private variants is avoided.

The complexity of each of the major hgs, as seen from the M-J-networks, Fig 4B-D, with multiple nodes and a span of many mutations between different leaves suggests that the hgs are not representative of single early founder event, as one mtDNA mutation is expected to occur per 3,000 yrs^66^. A notable exception is hg A, with a total prevalence of ~ 0.5 %, where the largest node, figure 4B, is associated with “daughter” nodes only 1-2 mtDNA mutations from the major node. It was only possible to identify the sub-hgs A1 and A5, and they only constituted a small fraction of the total A hg. A likely source of the A hgs is the Inuit population from Greenland, that is now a self-ruling part of the Kingdom of Denmark. Several studies have established that the major hgs in Greenlandic Inuits is A2 followed by hg D3 ^67-70^, whereas other Inuit populations have other characteristic hgs^70; 71^.

The lack of recombination, high mutation frequency, and fixation of mtDNA hgs has enabled the use of mtDNA in population genetics to study population ancestry, migrations, gene flow and genetic structure^72^; ^73^. Thus, populations on different continents, e.g. Native Americans^74^, Africans^75^ and Europeans^76^ were ascribed specific matrilineal mtDNA hg distributions^62^; ^77^. However, recent studies using autosomal SNP markers have disclosed a considerable ancestral complexity underlying an mtDNA classification in specific admixed populations, and the prediction of a specific mtDNA hgs is not possible from a specific continental ancestry based on nuclear genetic markers^78^. A problem with genetic analysis of admixed populations is the lack of temporal resolution. It is often not possible to date a specific population split because differentiation between new and old migrations is impossible. Recent advances in sequencing ancient genomes^79^ have made it possible to combine genetic information from ancient humans with archaeological information on the age of skeletal remains^80^, thus constructing hg distribution maps with a temporal dimension for e.g. Ice age and Bronze age^81^; ^82^ Europe. Maps, that may help explain demic and cultural exchange^70^; ^83-85^.

There is no solid evidence of the presence of humans in Denmark ^86^ ^87^ prior to the Last Glacial Maximum (LGM)^88^ (26.5 – 19 kYBP). At that time Denmark was covered in ice, except for the south-western part of Jutland^89^, and following the retraction of the ice sheath, peopling became possible from the south^89-92^. The first inhabitants documented were late Paleolithic hunters entering from southern Europe^93^. These hunters are discernible from Bølling time^89^ around 12,800 BC. The archaeological remains suggest their transient presence in seasonal hunting periods^94^ until the Mesolithic around 9,700 BC. The Hamburgian, Federmesser, Bromme and Ahrensburgian material cultures^89^; ^95^, well known from findings in Germany^89^, are represented. The earliest anatomically normal humans (ANM), present from around 45 – 41.5 kYBP^96-98^, where ancient DNA studies have revealed the presence of mtDNA hgs M and pre-U2^99^, have little similarity to present-day Europeans. However the Europeans from around 37 kYBP to 14 kYBP have left their mark on present-day Europeans^82^. From 14 kYBP the European population has a strong near-eastern component^82^. Around 7 kYBP the Neolithic transformation gradually started as the result of a demic dissemination of Neolithic Aegeans^100^. Thus, a minimum of three ancestral populations, i.e. a western European hunter-gatherer, an ancient north Eurasian, and an early European farmer population are needed to explain present-day European autosomal genome compositions ^101^. In Denmark, where Paleolithic ancient genetic data have not been published, ancient mtDNA haplotyping of three Neolithic corpses from 4.2 – 4 kYBP revealed two U4 and one U5a mtDNA hgs^102^ and later samples showed a mixture of hgs also found in northern Germany^99^; ^102^. The presence of mtDNA clades deriving from Paleolithic and Neolithic Europeans in the extant Danish population is thus explained.

In the Bronze age, an influx of people from the Russian steppe and North Caucasus, bringing the Indo-European language and culture^103^, resulted in the last major prehistoric demic change^81^ in Europe. In historic times the migrations have been many, particularly in the half-millennium following the fall of the Western Roman Empire^104^. In Denmark, apart from continuous demic exchange with Southern Scandinavia and present-day Germany^89^; ^92^, early historic time was mostly characterized by emigration, i.e. the Heruli, Cimbriae and Teutones, Burgundians and Vikings^91^; ^92^. The first census held in 1769 AD reported the population of what is present day Denmark to be ca 800,000 persons^105^. This number has increased to 5.6 mio in 2014, despite emigration of 287,000 persons between 1867 and 1914 (~10% of the population) largely to Northern America, and the same number from 1914 to 1968^106^. After the Second World War, Denmark has seen a considerable immigration from other European countries, but also from Asian, Middle Eastern and African countries, Suppl Figure 3. This part of Danish history can explain the occurrence of a diversity of mtDNA clades deriving from a large number of geographic regions. The extant distribution of mtDNA hgs reported here, Tables 1-4, reflect the complicated and heterogeneous origins of the people currently inhabiting Denmark.

The distribution of H sub-hgs, Table 3, and the M-J graph of the H-hg, Figure 4A, with multiple major nodes and 5-10 variants between leaves, suggest that the H-hg lineages are the result of repeated immigrations. Most likely from northern Europe, where a study of 39 prehistoric hg H samples has shown that the distribution of hg H differed between early and middle-to-late Neolithic groupings. Prior to Neolithicum H hgs have not been found in skeletal remains; an ensemble of Swedish Mesolithic hunter-gatherers all had U-hgs^83^. Whereas H, H5 and H1 were found throughout Neolithicum, H5b, H10, H16, H23, H26, H46, H88, and H89 were seen in early Neolithic samples, and H2, H3, H4, H5a, H6, H7, H11, H13, H82, and H90 in middle-to-late Neolithic samples^107^. The extant Danish H-hg distribution is compatible with contributions from throughout Neolithicum, but estimating when the hgs appeared in Denmark remains unfeasible, as the distribution is similar to that of northern Germany. However, not all carriers of an H-hg have a Danish autosomal geno-geographic affinity, table 5, compatible with a considerable recent immigration from countries with a European mtDNA distribution, Suppl Figure 3.

The most frequent Danish U-sub-hg is U5, Table 3, which is an old European mtDNA hg, with two major subclades, U5a and U5b, with coalescence time estimates of 16 – 20 kYBP and 20 – 24 kYBP, respectively^108^. The U5-hg is the most frequent U-sub-hg after LGM^99^ and the carriers have a Danish autosomal genomic ancestry around 90 %, and a European ancestry of 6.1 – 8.8%, Table 5. However, several of the less frequent U-sub-hgs have a considerable, ~ 40 – 60 % non-Danish and non-European geno-geographic affinity based on autosomal markers, with U7 related to South East Asia and U6 to the Middle East, respectively. In addition, J & T hgs, Figure 5, exhibit a relatively strong non-Danish autosomal genomic ancestry. The M-J graphs, Figure 4A, also disclose a considerable variation, much larger than could be attained merely within the timeframe where Denmark has been populated. This may be explained by an extensive immigration, and for U6 and U7 in very recent time. This is also compatible with the rising frequency of these hgs during the 25- year study period, Figure 6 and Suppl Table 3.

The N-macro-hg exhibits a high (>90%) combined Danish and European genomic affinity, Table 5, suggesting that the major part has been in Denmark for long. The major N-hgs are the I-hg, which has been found in meso- and neolithic Scandinavians^102^ and hg-X^109^ and hg-W, Table 4, and they exhibit, figure 4B, a very extensive heterogeneity. All three hgs are old and have a broad, low frequency, distribution in western Eurasia^61^, resulting from migratory events from the Near East and Central Asia. These events, viz. the significance for the presence of the hgs in Denmark, can not be temporally resolved.

Macro-hg-M, albeit infrequent (1.6 %), Table 1, has also increased in frequency recently, Figure 6, and exhibits extensive heterogeneity, Figure 4C, in the M-J graph, suggesting many migratory events of diverse origins. The largest group is M, that could not be decomposed further, but the second most frequent, D4 (14.2% of M-hgs), is most likely of central Asian origin^110^, and the M2, M3, M4 and M6 hgs, constituting 12.5% of M-hgs, are of Indian or Pakistani origin^111^. These findings are compatible with the high genomic affinity (42.2%, Table 5) towards Central South Asia, and a recent entry to Denmark. There is a propensity for location in Metropolitan areas, Suppl Table 2, and the Capital region, Table 6, also compatible with recent immigration.

The low, but increasing, frequency of L-hgs (Table 1, Table 7 and Figure 6) predominantly L2 and L3 (Table 1), is also the result of multiple immigration events, as the M-J plot reveals an extensive heterogeneity (Figure 4D). L2 and L3 hgs are frequent in south, west or east Africa, whereas the contribution from central Africa is small^112^; ^113^. However, the admixture analysis, Table 5, suggests a much stronger autosomal genomic affinity to the Middle East (70.8 %) than to Africa (8.4%). However, this finding could be an artefact of the ADMIXTURE analysis, as the highly variable African genomic ancestry^59^ ^114^ is represented by only 127 genomes (~ 6 % of the total reference set – see Materials and Methods), as compared to 178 genomes from the Middle East, making the definition of African ancestry imprecise. Alternatively, it could be caused by the extensive demic exchange that have occurred through time between the Near East and Northern Africa^115^; ^116^. The number of children born in the period 1980 to 2005 with one or two African parents, See Suppl Figure 3, is compatible with the origin of the L hg persons being African. The L hgs are predominantly located in the Capital Region, Table 6, and in Metropolitan areas, Suppl Table 4, which is also compatible with recent immigration.

Immigration to Denmark increased from 1980, where 135,000 immigrants were registered, to 2005, where this number had risen to 345,000^105^. In the same period the number of descendants of immigrants, i.e. persons that might turn up in this study, rose from 18,000 to 109,000. Roughly 50 % of the immigrants were from western countries^105^, See Suppl Figure 3. This should give approximately 45,000 descendants of non-western immigrants over the time of study. This is within the order of magnitude to be expected from the frequency distribution of mtDNA hgs in Denmark. There is thus a reasonable concordance between the suspected number of immigrants, from the temporal and structural study of mtDNA hgs, and the registered births of persons with non-Danish parents.

Whereas the temporal change in mtDNA distribution can be explained by immigration, it is more difficult to explain the spatial clines, Table 6 and Suppl Table 2. The frequency of hg-H is higher in Northern Jutland than in other regions, particularly the Capital Region, and it is also higher in rural areas (p = 0.0001), Suppl Table 2. The difference cannot be explained by the comparatively much smaller differences in frequencies of hgs L and M. As the mobility of Danes was fairly restricted until around 1900^91^, it may represent differences caused by centuries of relative isolation of a population north of Limfjorden. In Slovenia historically confirmed geographically-based sub-stratification of the population^117^ has lead to extreme differences in mtDNA distributions.

A recent fine-scale gDNA population structure study from the UK^46^ revealed considerable geographical heterogeneity and enabled the identification of specific sources of admixture from continental Europe. A gDNA study of admixture in the Danish population^118^ showed considerably more homogeneity, but medieval admixture from Slavic tribes in North Germany, as well as a North-South gradient were discernible. Both of these studies limited the participants to persons with local grand-parents, whereas our present study was not directed towards previous generations, but rather present-time and prospective. In addition to the gDNA/mtDNA interaction, the demonstrated spatio-temporal dynamics of the mt DNA hg distribution should be taken into account when designing studies of mtDNA associations with disease, physiological characteristics and phamacological effects. A prerequisite for genetic association studies is that the individuals are sampled from a homogenous population, or that cryptic population structure is corrected for, if not, false positive associations are likely^119^ ^120^; ^121^.

Thus, when studying bi-genomic, i.e. both nucleo- and mito-genomic, disease associations, our results indicate that it is necessary to compensate for gDNA and mtDNA genetic stratification, as well as the interaction between the two sources of variation and spatio-temporal clines. To our knowledge, this has never been done, suggesting that previous reports on disease associations and mtDNA haplogroups should be considered preliminary.

## Acknowledgements

The iPSYCH study was funded by The Lundbeck Foundation Initiative for Integrative Psychiatric Research. We further gratefully acknowledge the financial support of The Jascha Foundation, The Strategic Research Council (“Heart Safe”), The Augustinus Foundation, The Jascha Foundation, The Lundbeck Foundation (Grant no. R67-A6552), and Familien Hede Nielsens Fond. This research has been conducted using the Danish National Biobank resource, supported by the Novo Nordisk Foundation.

